# Evaluating the efficacy of commercially available antisense oligonucleotides to reduce mouse and human tau *in vivo*

**DOI:** 10.1101/2022.12.29.522258

**Authors:** Pranav Vemula, Kathleen M. Schoch, Timothy M. Miller

## Abstract

Tauopathies, including Alzheimer’s disease (AD), are neurodegenerative diseases characterized by the accumulation of tau protein encoded by the *MAPT* (Microtubule Associated Protein Tau) gene. Various strategies targeting mechanisms to reduce tau pathology have been proposed and several tau-directed therapies are being investigated in clinical trials. Our lab previously developed a novel strategy to lower tau protein levels using antisense oligonucleotides (ASOs), showing that human tau (hTau) reduction in aged PS19 tauopathy mice reversed phosphorylated tau pathology, spared neurons, and prolonged survival. Currently, the tau-lowering ASO is being evaluated in the clinical trials with successful phase 1b results. Similarly, preclinical and clinical studies have demonstrated the use of other ASOs as effective therapeutic strategies. Acquiring ASOs for research purposes may be limited by partnerships with pharmaceutical companies. However, ASOs can be obtained through commercial vendors. The current study evaluates the efficacy of mouse and human tau-targeting ASOs obtained from a commercial vendor in various mouse models. We show that mice treated with purchased ASOs distribute among various brain cell types including neurons, microglia, and astrocytes. Mice treated with tau lowering ASOs show decreased mouse or human tau mRNA and protein levels. In addition, human tau lowering ASO-treated PS19 mice showed decreased phosphorylated tau (AT8) and gliosis relative to saline-treated PS19 mice. The results obtained in PS19 mice are consistent with data obtained from our previous study using a non-commercial tau-lowering ASO. Overall, the present study demonstrates the efficacy of commercially-available tau targeting ASOs *in vivo* to support their broad use by researchers.

## Introduction

A key shared feature of neurodegenerative diseases is the aberrant expression or accumulation of dysfunctional proteins. Tauopathies, including Alzheimer’s disease (AD), are the most common neurodegenerative diseases characterized by the abnormal aggregation of the microtubule associated protein tau (MAPT) in the brain (Götz et al., 2019). Various strategies targeting mechanisms to reduce tau pathology have been proposed and several tau-directed therapies are being investigated in clinical trials (Congdon & Sigurdsson, 2018; Jadhav et al., 2019; Guo et al., 2022). Unfortunately, clinical trials of tau-targeting or other agents for the treatment of tauopathies have shown limited progress. Thus, there is a need to pursue alternative strategies or disease-modifying therapeutics to reduce or prevent tau pathology. Gene targeting strategies, like antisense oligonucleotides (ASOs) that target mRNA to alter the protein expression, could be a viable approach to investigate disease mechanisms and test tau-targeted therapeutics that can ameliorate disease.

ASOs are single-stranded DNA sequences composed of a phosphate backbone and sugar rings, 8-50 base pairs in length, that bind complementarily via Watson-Crick-Franklin base pairing to the target RNA (Schoch & Miller, 2017). Unmodified ASOs are prone to degradation; therefore, chemical modifications made to the phosphate backbone and sugar rings decrease or prevent degradation by nucleases and enhance target affinity (DeVos & Miller, 2013a; Schoch & Miller, 2017). Upon binding to target RNA, the resulting ASO:RNA pairing recruits the enzyme RNaseH to degrade the mRNA transcript. This strategy has been used to lower the production of disease-causing genes/proteins in several diseases including amyotrophic lateral sclerosis (Smith et al., 2006; Jiang et al., 2016; Boros et al., 2022), Huntington’s disease (Kordasiewicz et al., 2012; Lane et al., 2018; Kingwell, 2021), and spinal muscular atrophy (Wurster & Ludolph, 2018).

Tau-targeted ASOs are currently being tested in clinical trials for the treatment of tauopathies, including Alzheimer’s disease (Mignon et al., 2018). In preclinical testing, this human tau lowering ASO reduced phosphorylated tau deposition, rescued hippocampal loss, and increased survival in a tauopathy mouse model (DeVos et al., 2017). Intrathecal administrations of IONIS-MAPTRx (a second generation 2’-O-methoxyethyl chimeric ASO designed to reduce tau expression) in non-human primates resulted in a mean *MAPT* mRNA reduction of 77% in the frontal cortex and 74% in the hippocampus without side effects (DeVos et al., 2017). These findings led to the design of the first-in-human study of human tau lowering ASO, IONIS-MAPTRX (Mignon et al., 2018). Currently, human tau lowering ASOs are continuing through clinical trial testing with a recent successful phase 1b study having met its primary objective of safety and tolerability. In this phase 1b trial, a dose-dependent reduction in the concentration of total tau in the CSF was observed in the patients receiving tau lowering ASO relative to the placebo group (Mummery et al., 2021). Together, these studies show that direct tau mRNA targeting can be an efficient and promising therapeutic strategy.

Given their experimental and clinical utility, ASOs are considered powerful research tools to inform upon disease mechanisms prompting many researchers to use ASOs in their research. However, the design and synthesis of ASOs often requires partnerships with pharmaceutical industries which may be not feasible for every researcher. Several companies can also create ASO reagents, but these must be tested by the end-user. Here, we obtained commercially available ASOs that lower mouse or human tau mRNA levels and tested their efficacy in various mouse models. Both mouse tau lowering and human tau lowering ASOs significantly reduced their target mRNA and protein. When tested in the PS19 mouse model of tauopathy, the human tau lowering ASO was also effective at ameliorating phosphorylated tau pathology and gliosis, consistent with previous studies. Commercially available tau-targeted ASOs effectively reduce mouse or human tau *in vivo*, establishing ASO reagents as tools for tau-directed studies.

## Methods

### Mice

Male C57BL/6 mice were purchased from Jackson Laboratories (stock number 000664) and used for ASO treatment studies at 4 months of age. hTau transgenic mice (Andorfer et al., 2003) and PS19 transgenic mice (Yoshiyama et al., 2007) were bred in-house and identified for transgene expression as previously described (Schoch et al., 2016; DeVos et al., 2017). For ASO treatment studies, we used male and female hTau mice at approximately 9 months of age and male and female PS19 mice and non-transgenic littermates at approximately 6 months of age. Animals were maintained on a 12-hour light/dark cycle and provided *ad libitum* access to food and water. All husbandry and experimental procedures were approved by the Institutional Animal Care and Use Committee at Washington University in St Louis.

### Antisense oligonucleotides

Antisense oligonucleotides (ASOs) were purchased from Integrated DNA Technologies (IDT, Coralville, Iowa, USA) designed with 2’-O-methoxyethyl (2’-MOE) sugar ring and phosphorothioate (PS) backbone modifications. The 2’-MOE modification is well-tolerated, enhances target affinity, and increases resistance to degradation by nucleases while the PS modification enhances stability and cellular uptake (Schoch & Miller, 2017). Tau lowering ASOs were designed using a “gapmer” strategy, which consists of 2’-MOE modifications flanking a central region of unmodified nucleotides. A control ASO with same modifications but without a specific target was used in select experiments. ASO names and sequences are listed in Table 1. Control and tau-targeting sequences were identical in both sequence and chemistry to previously published studies in C57BL/6 (DeVos et al., 2013), hTau (Self et al., 2018), and PS19 mice (DeVos et al., 2017).

**Table 1.** Commercially available control and tau-targeting ASOs used to evaluate tau lowering in mice. The species target and the mouse model in which the ASO was used are indicated. ASO modifications are indicated as follows: 2’-MOE nucleotides (red font), unmodified nucleotides (underlined). Nucleotides with PS backbone are indicated by asterisk. Abbreviations: mTau, mouse tau; hTau, human tau; KD, knockdown.

| ASO name | Target | Used in | ASO sequence (5' to 3') |
| --- | --- | --- | --- |
| mTau Control | No target | C57BL/6 mice | C*C*T*T*C*C*C*T*G*A*A*G*G*T*T*C*C*T*C*C |
| hTau Control | No target | hTau mice | C*CTAT*A*G*G*A*C*T*A*T*C*C*AGG*A*A |
| mTau KD | Mouse tau | C57BL/6 mice | A*T*C*A*C*T*G*A*T*T*T*T*G*A*A*G*T*C*C*C |
| hTau KD 1 | Human tau | hTau mice | C*CGTT*T*T*C*T*T*A*C*C*ACC*C*T |
| hTau KD 2 | Human tau | PS19 mice | G*C*T*T*T*T*A*C*T*G*A*C*C*A*T*G*C*G*A*G |

### Surgical procedures, euthanasia, and tissue dissection

C57BL/6, hTau, and PS19 mice were anesthetized with isoflurane via continuous inhalation and placed into a stereotaxic frame during surgical procedures. For ASO infusion into the lateral ventricle of C57BL/6 and PS19 mice, 28-day ALZET osmotic pumps (Durect, Model 2004) were implanted as previously described (DeVos & Miller, 2013b). hTau mice received a single intracerebroventricular (ICV) bolus injection as previously described (DeVos & Miller, 2013b). For euthanasia and tissue collection, mice were anesthetized with isoflurane and perfused with ice-cold 1X phosphate buffered saline (PBS). Brains were collected rapidly, and the left hemisphere (contralateral to the catheter placement or ICV injection) was drop-fixed into ice-cold 4% paraformaldehyde for histological processing. The right hemisphere (ipsilateral to the catheter placement or ICV injection) was dissected into separate pieces for RNA and protein analyses, snap frozen in liquid nitrogen, and stored at -80°C until use.

### RNA isolation and quantitative RT-PCR analysis

Total RNA was extracted from right hemisphere brain lysates using a QIAGEN RNeasy kit (Catalogue #74104, QIAGEN, Hilden, Germany) per manufacturer’s instructions. Gene targets were amplified using Express One-Step Superscript qRT-PCR universal kit (Catalogue #11781200, ThermoFisher Scientific) and measured on a QuantStudio 12K Flex Real-Time PCR system (ThermoFisher Scientific). Target mRNA expression levels were normalized to mouse *Gapdh* mRNA levels and compared using the ΔΔCt method. Primer and probe sequences were as follows: mouse total *Mapt*: forward 5′-GAACCACCAAAATCCGGAGA-3′, reverse 5′-CTCTTACTAGCTGATGGTGAC-3′, probe 5′-/56-FAM/CCAAGAAGGTGGCAGTGGTCC/3IABkFQ/-3′; human total *Mapt*: forward 5′-AGAAGCAGGCATTGGAGAC-3′, reverse 5′-TCTTCGTTTTACCATCAGCC-3′, probe 5′-/56-FAM/ACGGGACTGGAAGCGATGACAAAA/3IABkFQ/-3′; *Gapdh*: forward 5′-TGCCCCCATGTTGTGATG-3′, reverse 5′-TGTGGTCATGAGCCCTTCC-3′, probe 5′-/56-FAM/AATGCATCCTGCACCACCAACTGCTT/3IABkFQ/-3′.

### Protein homogenization and enzyme-linked immunosorbent assay (ELISA)

Total protein was extracted from tissue sections from the right hemisphere of mouse brain using RAB buffer (100mM MES, 1mM EDTA, 0.5mM MgSO4, 750mM NaCl, 20mM NaF, 1mM Na3VO4) containing phosphatase and protease inhibitors and homogenized with a handheld tissue homogenizer. Bicinchoninic Acid (BCA) assay was used to quantify extracted protein.

For quantification of human and mouse tau protein, sandwich ELISA of tau-5 coating antibody (20 µg/ml; Millipore Cat# 577801-100UG RRID: AB_212534) with either biotin-conjugated HT-7 (for human tau, 0.3 μg/ml; ThermoFisher Scientific Cat# MN1000B RRID: AB_223453) or BT-2 (for mouse tau, 0.3 μg/ml; ThermoFisher Scientific Cat# MN1010B RRID: AB_10974155) antibodies was performed. Initially, 96-half-well plates were coated and incubated overnight with tau-5 antibody diluted in carbonate coating buffer (0.2M NaHCO3/Na2CO3). The plates were then blocked with 4% bovine serum albumin (BSA)/PBS at 37°C. Standards of recombinant human tau (2N4R; rPeptide, Bogart, GA) or mouse tau (mTau40, 432aa, produced in the laboratory of Eva-Maria Mandelkow) and samples were diluted in sample buffer (0.25% BSA, 300mM Tris, and 1X protease inhibitor cocktail in PBS), loaded on plates, and incubated overnight at 4°C. HT7 and BT-2 capture antibodies followed by streptavidin-poly-HRP-40 conjugate (1:4000; Fitzgerald, Acton, MA) were applied and detected by the addition of 3,3’,5,5’-tetramethylbenzidine liquid substrate, Super Slow (T5569, Sigma) reagent. The plates were read using a BioTek microplate reader (Biotek Epoch, Winooski, VT) at 650nM.

### Immunohistochemistry

The left hemisphere of mouse brains remained in 4% paraformaldehyde for 24 hours, was transferred to a 30% sucrose solution for cryoprotection, and frozen in cold (−20°C to -35°C) 2-methylbutane. Frozen brains were sectioned into 40 µm coronal sections using a microtome and stored free-floating in a cryoprotectant solution (30% ethylene glycol, 30% glycerol, 10% 0.2M phosphate buffer, in water) at -20°C until use.

For immunofluorescence, three sections per brain (approximate bregma level range - 1.95mm to -2.98mm) were selected from saline- and ASO-treated PS19 mice and non-transgenic littermates. All immunofluorescence procedures were performed at room temperature except primary antibody incubation. Sections were incubated in a Tris-buffered saline (TBS)-0.1% Triton X-100 blocking solution containing 2% normal donkey serum, 4% BSA, 0.1% hydrogen peroxide, and 100mM glycine and then incubated in primary antibody at 4°C overnight (Pan ASO 1:1000, gift from Ionis Pharmaceuticals, Carlsbad, CA, USA; NeuN 1:500, 266004 Synaptic Systems, Goettingen, Germany; GFAP 1:1000, AB5541 Millipore-Sigma; Iba1 1:500 ab5076 Abcam, Cambridge, United Kingdom). Sections were incubated in blocking solution prior to incubation in secondary antibodies (all at 1:1000; for ASO: anti-rabbit Alexa Fluor Plus 647, A32795 ThermoFisher Scientific; for NeuN: anti-guinea pig Alexa Fluor 555, A21435 Invitrogen or anti-chicken Alexa Fluor 594, 703-585-155 Jackson Immuno Research Labs, West Grove, PA, USA; for GFAP: anti-chicken Alexa Fluor 488, 703-545-155 Jackson Immuno Research Labs; for Iba1: anti-goat Alexa Fluor 488, 705-545-147 Jackson Immuno Research Labs). Sections were washed, incubated in DAPI (1:1000), and mounted on slides. Slides were covered with coverslips using Fluoromount-G mounting media (SouthernBiotech, Birmingham, AL, USA) and sealed with nail polish. To visualize cell type-specific ASO localization, z-stack images (30µm depth with 3µm step size) were captured at 20x on a Nikon A1Rsi confocal microscope (Nikon, Minato City, Tokyo, Japan). To visualize overall microglial and astrocytic reactivity, slides were imaged at 10x on a Zeiss Axio Scan.Z1 slide scanner (ZEISS, Oberkochen, Germany).

For immunohistochemistry of phosphorylated tau, six sections per brain (approximate bregma level range -1.65mm to -2.98mm) were selected from non-transgenic and saline-or ASO-treated PS19 mouse tissue. All immunohistochemical procedures were performed at the room temperature except primary antibody incubation. Sections were incubated in 0.3% hydrogen peroxide prior to blocking in 3% nonfat dry milk in PBS-0.25% Triton X-100 solution. Sections were then incubated in primary antibody at 4°C overnight (biotinylated AT8, 1:500, MN1020B, ThermoFisher Scientific). The biotin signal was amplified using an ABC Elite kit (1:400; PK-6100, Vector Laboratories, Burlingame, CA, USA) and detected by reaction with 3, 3’-diaminobenzidine (DAB) solution (SK-4100, Vector Laboratories). Sections were then washed and mounted on slides. Mounted sections were dehydrated through an ethanol gradient followed by incubation in xylene. Slides were covered with coverslips using mounting media (Cytoseal 60, #8310-4, ThermoFisher Scientific) and imaged at 20x on a Hamamatsu NanoZoomer HT whole slide imager (Hamamatsu Photonics, Japan).

### Immunohistochemical image analysis

To quantify phosphorylated tau, section images from AT8 immunohistochemistry were exported at 2.5x (cortex) or 5x (hippocampus) magnification using the NDP.view (Hamamatsu) program and analyzed in ImageJ as previously described (Schoch et al., 2016). In brief, cortical and hippocampal regions across six sections were outlined as ROIs and a uniform, global threshold used to identify AT8+ reactivity within each region. The amount of AT8+ reactivity was recorded as a percentage of the total ROI area.

To quantify gliosis, images from saline-and ASO-treated PS19 and non-transgenic littermate controls stained for Iba1 (microglia) and GFAP (astrocytes) were exported using ZEN 3.4 software (ZEISS) and analyzed in ImageJ. Hippocampal regions across three sections were outlined as ROIs and a uniform, global threshold used to identify Iba1+ or GFAP+ reactivity within each region. The amount of reactivity was recorded as a percentage of the total ROI area.

### Statistical analysis

Data were represented in mean ± SEM and analyzed with GraphPad Prism (version 9). mRNA levels and protein expression were analyzed using unpaired t-test (for two groups) or ordinary one-way ANOVA with Tukey’s post-hoc test (for three or more groups). Analysis of immunohistochemical data (i.e., phosphorylated tau and gliosis) was done using one-way ANOVA with Tukey’s post-hoc test.

## Results

### Commercial ASOs are taken up by neurons and glia

We have previously demonstrated that tau-targeted ASOs reliably distribute throughout the brain and can be taken up by neurons and glia in the central nervous system (CNS) (DeVos et al., 2013; DeVos et al., 2017). To test the ability of a commercial ASO to distribute and localize to cell types within the CNS, we performed immunofluorescence to visualize localization of the ASO in neurons, microglia, and astrocytes. We noted co-localization of the ASO with NeuN (Figure 1A-D, G-H), Iba1 (Figure 1A, E, G), and GFAP (Figure 1B, F, H), confirming neuronal and glial uptake of ASO.

**Figure 1:**
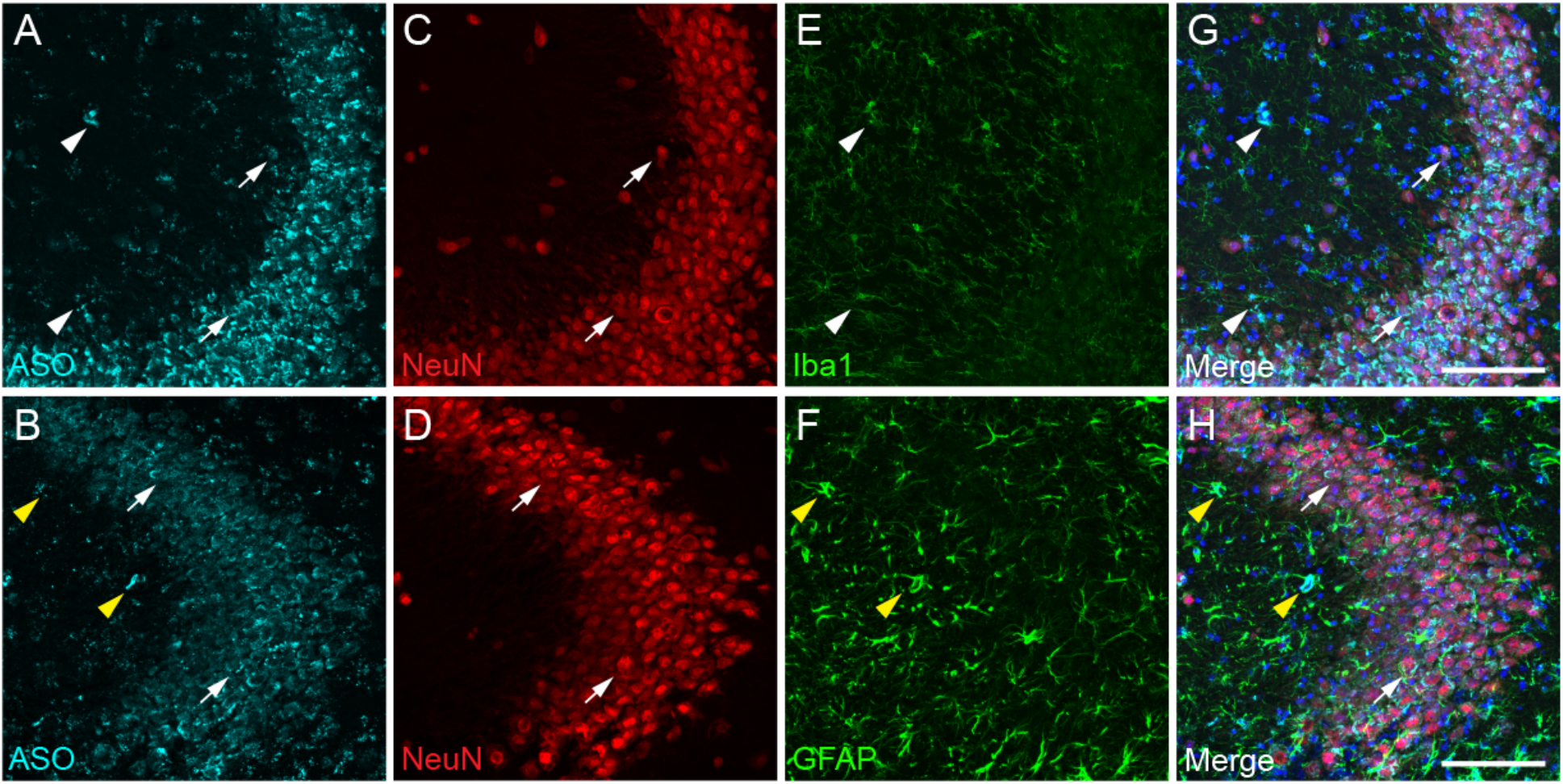
Commercially available ASOs distribute to neurons and glia within the brain. Immunofluorescent images within the CA3 region of the hippocampus from mice treated with purchased ASOs showing A-B) ASO distribution among C-D) neurons (NeuN), E) microglia (Iba1), and F) astrocytes (GFAP). G-H) Merged images demonstrate co-localization of the ASO with neurons (white arrows), microglia (white arrowheads), and astrocytes (yellow arrowheads). Scale bars = 100µm.

### Mouse tau knockdown in C57BL/6 mice reduces tau expression

Although mouse tau does not form pathological tau aggregates in wildtype mice, mouse tau may be used as a target to inform upon the physiological function of tau within the CNS. To test the effect of a commercially available mouse tau-targeted ASO, we administered mouse tau knockdown (mTau KD) ASO via intraventricular osmotic pump to C57BL/6 mice and measured mTau mRNA expression two months after pump implantation. C57BL/6 mice treated with mTau KD ASO showed significant reduction in the levels of mTau mRNA expression levels (Figure 2A; One-way ANOVA, F=74.68, P<0.0001; Tukey’s multiple comparisons test) relative to saline (73% decrease; P<0.0001) or control ASO (68% decrease; P<0.0001) treated mice. No significant difference in mTau mRNA expression was observed between saline and control ASO treated mice (P=0.1265). We also measured changes in the protein expression of mTau protein after ASO administration by ELISA. We found reduction in the levels of mTau protein in mice treated with mTau KD ASO (Figure 2B; One-way ANOVA, F=12.69, P<0.01; Tukey’s multiple comparisons test) compared to saline (61% decrease; P<0.01) and control ASO (56% decrease; P<0.01).

**Figure 2:**
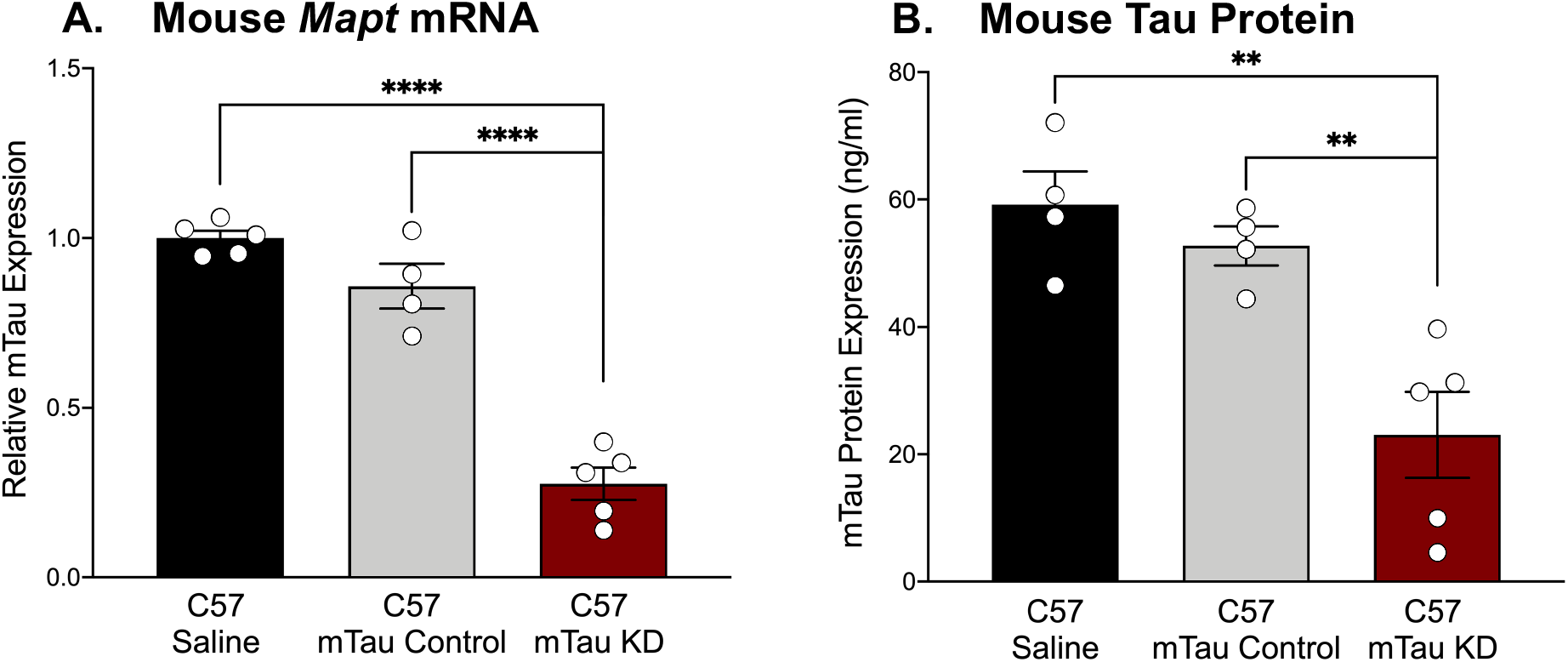
Purchased mouse tau-targeting ASO reagents reduce mouse tau mRNA and protein in C57BL/6 mice. A) Mouse tau mRNA expression was significantly decreased following administration of purchased mouse tau knockdown (mTau KD) ASO in C57BL/6 (C57) mice relative to saline and control ASO treated mice (n=4-5/group). Mouse tau mRNA levels were normalized to *Gapdh* and shown relative to saline treated mice. B) Mouse tau protein expression was significantly decreased in mTau KD ASO-treated mice relative to saline or control ASO-treated mice (n=4-5/group). Data are presented as mean±SEM. **P<0.01, ****P<0.0001.

### Human tau knockdown in hTau mice reduces human tau mRNA and protein expression

Human tau mice (hTau mice) express all six isoforms of human tau under control of the human tau promoter in the absence of endogenous mouse tau (Andorfer et al., 2003), offering a model that closely mirrors human tau expression. Our previous study in hTau mice treated with hTau KD ASO showed significant reduction in human tau mRNA and protein levels (Self et al., 2018). To test the effect of the commercially available hTau KD ASO, we administered hTau KD ASO 1 via single ICV injection to hTau mice and measured hTau mRNA and protein levels. hTau mice treated with hTau KD ASO 1 afforded a significant reduction in the levels of human tau mRNA expression levels (Figure 3A; Two-tailed t-test, P<0.0001) relative to control ASO (84% decrease) treated mice. Using ELISA, we also measured human tau protein expression after hTau KD ASO 1 administration in hTau mice. ASO-mediated hTau knockdown in hTau mice showed a significant reduction in human tau protein expression relative to control ASO treated hTau mice (Figure 3B; 62% decrease, Two-tailed t-test, P<0.05).

**Figure 3:**
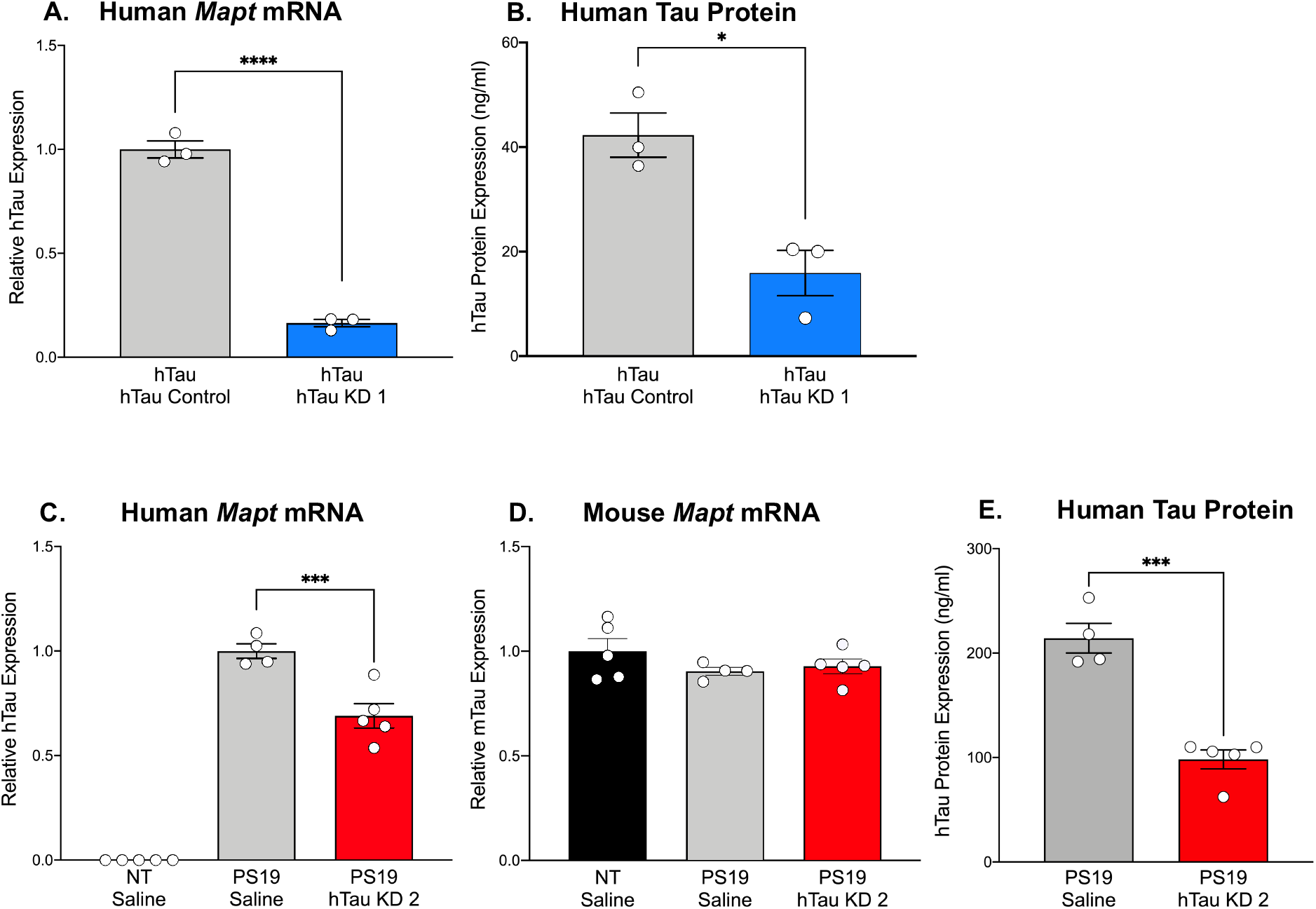
Purchased human tau-targeting ASO reagents reduce human tau mRNA and protein in hTau and PS19 mice. Human tau mRNA and B) human tau protein expression were significantly decreased following purchased human tau knockdown (hTau KD 1) ASO administration in hTau mice relative to control ASO treated hTau mice (n=3/group). Human tau mRNA levels were normalized to *Gapdh* and shown relative to control ASO. C) Human tau mRNA expression was significantly decreased following treatment with purchased human tau knockdown (hTau KD 2) ASO in PS19 mice relative to saline treatment (n=4-5/group). Human tau mRNA expression was undetected in non-transgenic (NT) littermates treated with saline (n=5). D) No difference in mouse tau mRNA levels was observed among treated mice. mRNA levels were normalized to *Gapdh* and expressed relative to saline-treated mice. E) Human tau protein expression was significantly reduced in hTau KD 2 ASO-treated PS19 mice relative to saline-treated PS19 mice. Data are presented as mean±SEM. *P<0.05, ***P<0.001, ****P<0.0001.

### Human tau knockdown in PS19 mice reduces human tau mRNA and attenuates pathology and gliosis

Mutant tau expression in PS19 closely recapitulates the time course of tau phosphorylation and aggregation, neuroinflammation, and cell loss evident in human tauopathies (Yoshiyama et al., 2007). In our past studies, we have demonstrated the ability of hTau-lowering ASOs to reverse tau pathology and extend survival in these mice (DeVos et al., 2017). Using a commercially available version of the ASO, we administered hTau KD ASO 2 via osmotic pump to PS19 mice and measured human tau mRNA and protein expression two months after ASO administration. Non-transgenic littermates treated with saline were included as treatment controls. hTau KD ASO 2 treated mice showed a significant reduction in the levels of human tau mRNA (Figure 3C; One-way ANOVA, F=164.9, P<0.0001; Tukey’s multiple comparisons test) relative to saline (31% decrease; P<0.001) treated mice without a change in mouse tau mRNA expression, confirming the specificity of the human tau target (Figure 3D; One-way ANOVA, F=1.258, P=0.3221). We observed reduction in human tau protein expression in PS19 mice treated with hTau KD ASO relative to saline treated PS19 mice (Figure 3E; 54% decrease, Two-tailed t-test, P<0.001).

PS19 mice develop phosphorylated tau at 6 months of age, accompanied by significant microgliosis beginning at 3 months age and astrogliosis at 6 months age (Yoshiyama et al., 2007). To investigate pathological outcomes in PS19 mice after hTau lowering with a commercially available ASO, we performed immunohistochemistry using the AT8 antibody to identify phosphorylated tau reactivity within the cortex and hippocampus (Figure 4A-D). PS19 mice treated with hTau KD ASO 2 exhibited reduced phosphorylated tau staining compared to saline treated PS19 mice in both the cortex (49% decrease) and hippocampus (39% decrease), although not statistically significant (Figure 4E, Cortex: One-way ANOVA, F=10.50, P=0.0028, Tukey’s multiple comparison test, P=0.1032; Figure 4F, Hippocampus: One-way ANOVA, F=8.162, P=0.0067, Tukey’s multiple comparison test, P=0.3126). Next, to test the effect of hTau KD ASO 2 on microglial and astrocytic activation, we probed mouse brain sections with antibodies to Iba1 (Figure 5A-B) and GFAP (Figure 5D-E). PS19 mice treated with hTau KD ASO 2 showed significant reductions in Iba1 (81% decrease) and GFAP (45% decrease) relative to saline treated mice (Figure 5C, Iba1: One-way ANOVA, F=9.216, P<0.01, Tukey’s multiple comparison test, P<0.01; Figure 5F, GFAP, F=15.20, P<0.001; Tukey’s multiple comparison test, P<0.01).

**Figure 4:**
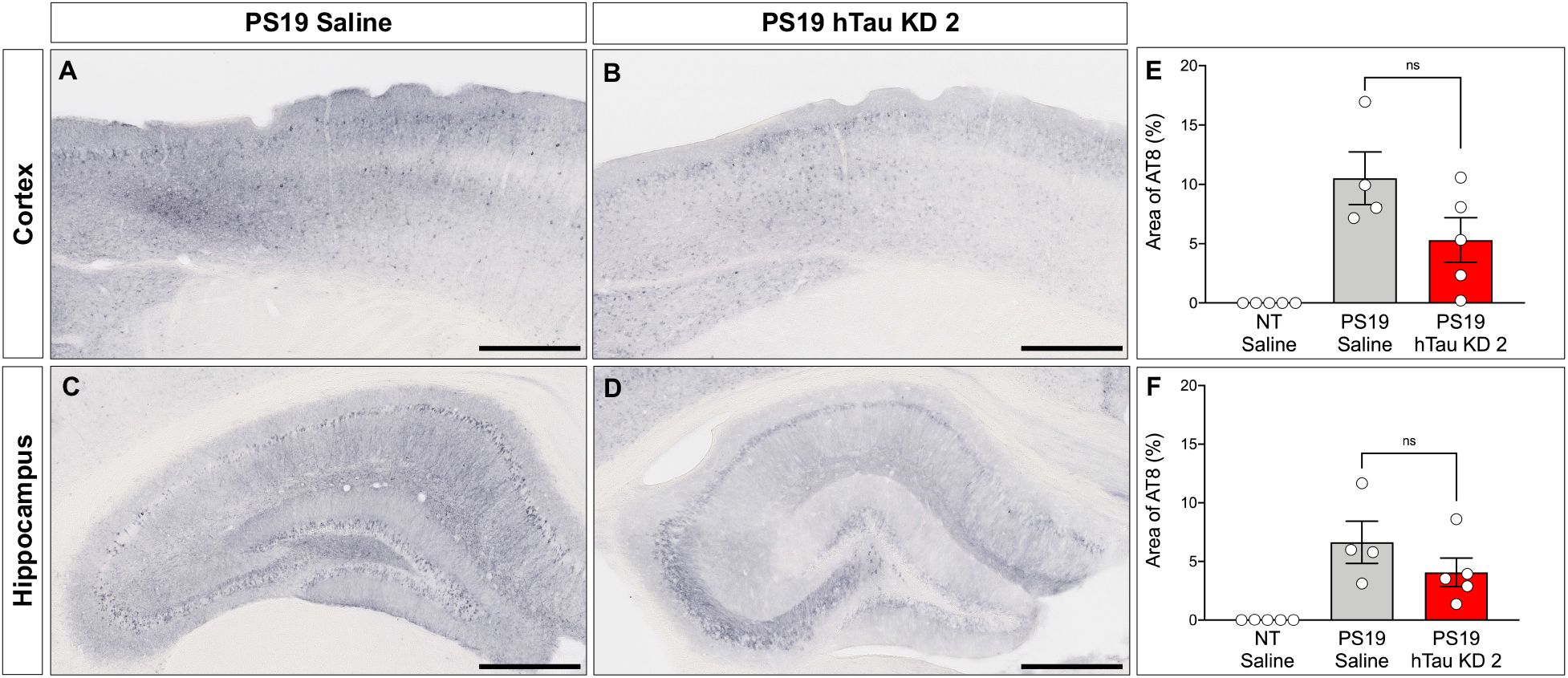
hTau KD ASO treatment in PS19 mice reduces phosphorylated tau accumulation. Representative images of phosphorylated tau (AT8) immunohistochemistry within the cortex and hippocampus from A-C) saline-treated PS19 mice and B-D) human tau knockdown (hTau KD 2) ASO-treated PS19 mice. Scale bar = 500µm. Quantification of the percentage of AT8 immunoreactivity within the E) cortex and F) hippocampus of non-transgenic (NT) and saline or hTau KD 2 treated PS19 mice (n=4-5/group). Data are presented as mean±SEM. ns, non-significant.

**Figure 5:**
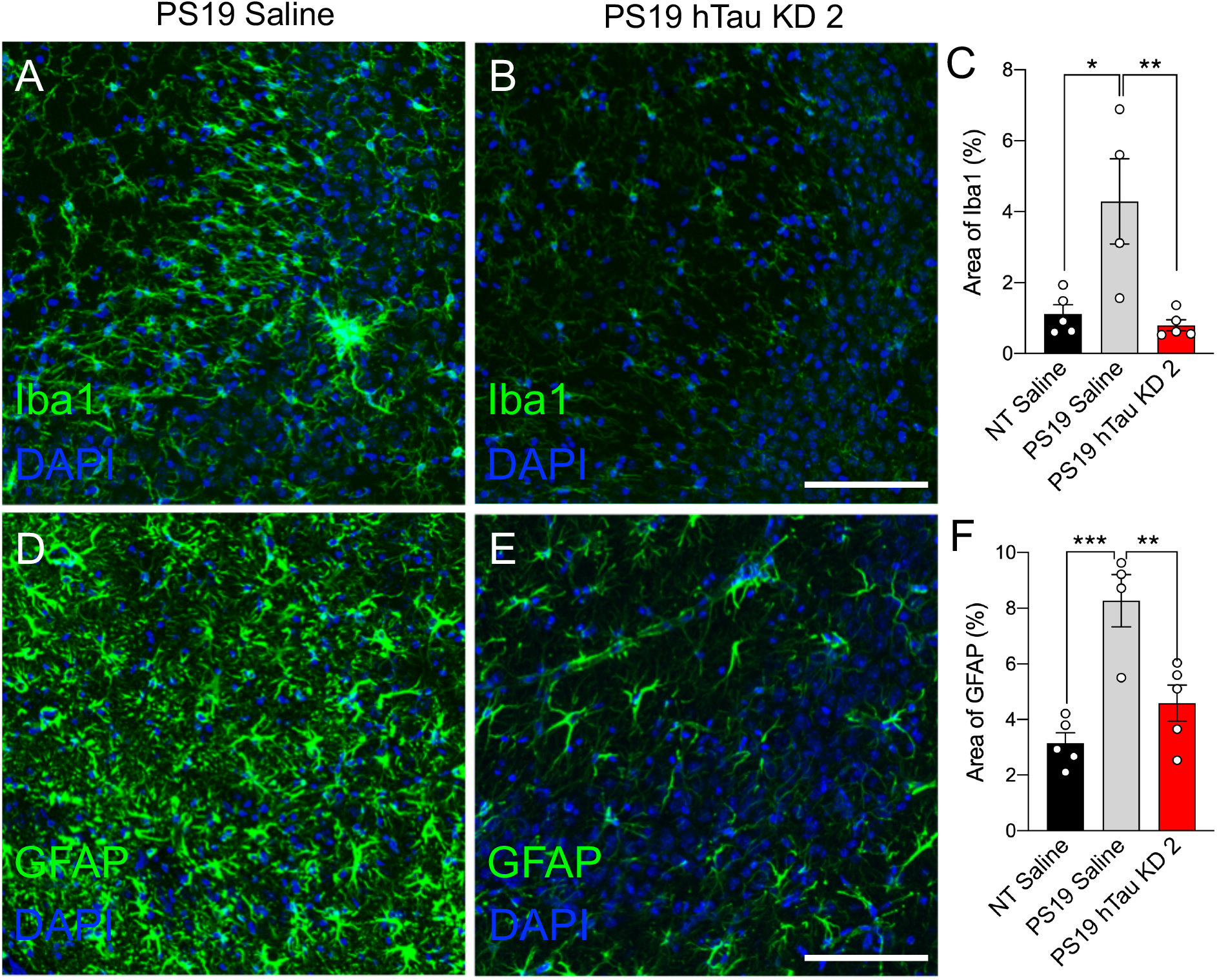
hTau KD ASO treatment in PS19 mice reduces gliosis. Representative images from the CA3 region of the hippocampus showing A-B) Iba1 and C-D) GFAP immunofluorescence for microglia and astrocytes, respectively, from PS19 mice treated with saline or human tau knockdown (hTau KD 2) ASO. Scale bars = 100µm. Quantification of the percentage of E) Iba1 and F) GFAP immunoreactivity within the hippocampus of non-transgenic (NT) and saline or hTau KD treated PS19 mice (n=4-5/group). Data are presented as mean±SEM. *P<0.05, **P<0.01 and ***P<0.001.

## Discussion

We tested commercially available mouse and human tau lowering ASOs in adult mice to support their use as research tools for tau-directed studies. We identified that the ASOs can be taken up by neurons and glia within the CNS, suggesting widespread distribution and uptake consistent with prior studies. Mouse- and human tau-targeted ASOs were able to reduce mouse or human tau mRNA and protein levels when administered to wildtype or human tau-expressing mice (hTau and PS19 mice), respectively. When administered to PS19 mice, hTau KD ASO also reduced phosphorylated tau deposition and gliosis.

Advancements in ASO chemistry coupled with our understandings of RNA biology have led to an evolution of ASO design and application (Crooke et al., 2021). Although the ASO sequences and modifications used in our studies were previously determined, designing ASOs to target a gene of interest has become feasible with freely available computational algorithms to generate potential ASO candidates for subsequent testing. Additionally, factors such as RNA secondary structure, GC content, and binding energy (ΔG°37) are considered when selecting potential ASOs for further screening (Chan et al., 2006). ASOs generated from an algorithm should be then chemically modified on both backbone and sugar rings to convert into a fully functional ASO. Among these modifications, the PS backbone modification is the most used as it renders ASOs with increased stability, protection from nucleases, increased binding ability, and the ability to recruit RNase H. For modification on sugar rings, fluoro, 2’-*O*-methoxyethyl (2’-MOE), 2’-*O*-Methyl (2’-*O*-Me), constrained ethyl (cEt), and locked nucleic acid (LNA) are commonly used. Particularly, 2’-MOE and 2’-*O*-Me modifications offer enhanced stability and a low pro-inflammatory response and are well-tolerated *in vivo* (Crooke et al., 2021). Chemical modification, synthesis, and purification of an ASO is a labor-intensive and needs expertise. However, ASO production can be outsourced to several companies for their ASO synthesis and purification services.

The commercially available ASOs used in the current study were synthesized and chemically modified using previously published *MAPT* targeting sequences (DeVos et al., 2013; DeVos et al., 2017; Self et al., 2018). The control and hTau KD ASOs used in the present study were administered into right lateral ventricle via ICV bolus injection for hTau mice and infused via osmotic pump implantation for C57 and PS19 mice. All ASOs tested effectively decreased *MAPT* suggesting commercially available ASO reagents are feasible for use. Previous studies have used tau targeting ASOs with alternative modifications showing similar efficacy. Sud et al. (2014) employed morpholino ASOs to target *MAPT* in the human neuroblastoma cell lines, SH-SY5Y and IMR32, and in tau transgenic mice. These ASOs reduced *MAPT* mRNA up to 50% and protein up to 80% (Sud et al., 2014). Another study demonstrated the ability of 2’-*O*-Methyl (2’-*O*-Me) modified ASOs on a PS backbone to reduce *MAPT* mRNA (92% reduction) and tau protein (50% reduction after 48 hour treatment) levels *in vitro* in SH-SY5Y cells (Chakravarthy et al., 2020). These studies underscore the capability of ASOs *in vitro* and *in vivo* as excellent reagents to lower tau levels.

Importantly, the tau lowering ASOs tested here can target mutant human tau, enabling studies on tau as a therapeutic target in tau-mediated diseases. DeVos et al. (2017) demonstrated that use of hTau-targeted ASOs could reverse tau pathology, preserve neuronal number, and increase survival in aged PS19 mice (DeVos et al., 2017). Consistent with this study, the commercially available equivalent ASO afforded reductions of phosphorylated tau in the cortex and hippocampus of PS19 mice relative to saline treatment. Although these reductions did not achieve statistical significance, we note the high degree of variability in AT8 reactivity within treatment groups, likely due to the inherent variability of the line and relatively small group sizes. In addition, both male and female PS19 mice were evaluated, and sex-dependent differences in phosphorylated tau reactivity have been reported (Sun et al., 2020). Although, gliosis was significantly attenuated in hTau KD ASO-treated PS19 mice, neurodegeneration and survival measures would further confirm the efficacy of the ASOs in a pathological context. However, the mice in the present study were evaluated at 6 months age, prior to expected neuronal loss or premature death. However, based on the ability of commercially available ASOs to attenuate gliosis, we speculate that these ASOs may be similarly efficient to attenuate late-stage pathology or other deficits, supporting their use as a therapeutic agent for researchers studying tau-mediated disease mechanisms.

To date, nine ASO drugs have been approved by the United States Food and Drug Administration for a variety of diseases (Dhuri et al., 2020; Crooke et al., 2021). With the development of nusinersen, an ASO indicated for spinal muscular atrophy, the use of ASOs in CNS disorders has continued to expand, and several ASOs targeting genes implicated in neurodegenerative diseases are currently in clinical trial. For example, tofersen, a 2’-MOE ASO designed to target superoxide dismutase (*SOD1*) for amyotrophic lateral sclerosis (ALS) (Miller et al., 2020; Miller et al., 2022) and BIIB080, an ASO that targets human *MAPT* gene to prevent tau protein production for tauopathies including AD (Mummery et al., 2021). On the other hand, there are ASOs, which were not effective in human clinical trials resulting in the clinical trial halt. For example, tominersen, a 2’-MOE ASO to target huntingtin for Huntington’s disease (ClinicalTrials.gov Identifier: NCT02519036; NCT03761849) (Tabrizi et al., 2019) showed no serious adverse effects in phase 2 trials, but phase 3 trials were stopped due to lack of efficacy. Similarly, a recent phase 1 clinical trial for C9orf72 ALS has been discontinued due to the lack of clinical benefit (Biogen, 2022). Regardless of these trial outcomes, preclinical testing of ASOs was vital to their successful transition into clinical application. ASO reagents that are accessible and effective may enable future therapeutic discoveries.

Alternative gene-targeting siRNA reagents or synthetic small molecules can serve as research tools or therapeutic drugs to achieve a similar effect to ASOs. siRNAs are synthetic 19-22 bp long double-stranded molecules that are complementary to target mRNA and can regulate mRNA expression (Watts & Corey, 2012). Like ASOs, siRNAs are advantageous because of their efficacy and unrestricted choice of targets. However, unlike ASOs, siRNAs often require a delivery agent such as liposomes or conjugation to cholesterol (de Fougerolles et al., 2007). For small molecules, which are synthetic organic compounds that target proteins, a major advantage is their low molecular weight, which enables cellular penetration. However, their development involves a complicated process of targeted validation approaches, they often exhibit short half-lives requiring frequent administration, and lack target specificity. While every drug modality has its advantages and limitations, target engagement is one of the most desirable drug properties, and ASOs, owing to their complementary base pairing, are highly specific. However, ASOs can elicit off-target toxicities and pro-inflammatory effects (Scoles & Pulst, 2018) and lack the ability to cross blood-brain barrier requiring direct administration into the CSF via a relatively invasive lumbar puncture procedure (DeVos & Miller, 2013a). Despite these limitations, the advantages of ASOs outweigh their alternatives and the ease of development and translatability make them ideal candidates as therapeutics (Schoch & Miller, 2017). Overall, the current study demonstrates the use and efficacy of commercially-available tau-targeting ASOs to show that easy access to ASOs through commercial vendors enables their broad use by researchers to address various research questions on disease mechanisms and potential therapeutics.

## Acknowledgements

This work was supported by Tau Consortium (T.M.M.). The microscopy work was supported by the Hope Center Alafi Neuroimaging Lab and an NIH Shared Instrumentation Grant award to Washington University (S10 RR027552), and performed in part through the use of Washington University Center for Cellular Imaging (WUCCI) supported by Washington University School of Medicine, The Children’s Discovery Institute of Washington University and St. Louis Children’s Hospital (CDI-CORE-2015-505 and CDI-CORE-2019-813) and the Foundation for Barnes-Jewish Hospital (3770 and 4642).

